# Deciphering the Soybean Rust Arsenal: Effectorome Studies and Host-Pathogen Interaction (HPI) network in *Phakopsora pachyrhizi* and *Glycine max* Pathosystem

**DOI:** 10.1101/2024.06.17.599355

**Authors:** E.S. Oliveira, W.A. Pereira

**Affiliations:** Universidade Federal de Lavras

**Keywords:** PPI, Pathogen Effector, Plant pathogen interaction, Effector triggered immunity

## Abstract

Asian soybean rust (ASR), cause by the fungal pathogen *Phakopsora pachyrhizi*, stands out as the paramount menace to soybean cultivation, resulting in staggering yield decreases that can surpass 90%. Despite extensive endeavors, the development of cultivars endowed with comprehensive or enduring resistance against the myriad pathotypes remains elusive, thereby underscoring the imperative for pioneering improvement tactics. Fungal effectors, orchestrating the infiltration and colonization of hosts, constitute pivotal determinants in the infection cascade, thus emerging as prime targets for unraveling pathogenicity mechanisms. In this investigation, we endeavored to integrate effectoromics analysis with host-pathogen protein-protein interaction networks (HPPI) to delineate the modus operandi of *P. pachyrhizi* effectors in subverting plant host immunity. By meticulously delineating the repertoire of predicted effectors and prognosticating protein-protein interactions between effectors and soybean target proteins, we discerned pivotal candidate targets. Notably, a repertoire of 324 host proteins was anticipated to engage in interactions with 21 effectors, predominantly hailing from the Lysophospholipase (LysoPLs) cohort. Particularly conspicuous was the identification of a robust effector candidate (PhapaK8108_7408047) implicated in the targeting of 171 soybean proteins. Its functional involvement in RNA splicing, chromatin organization, histone deacetylation, protein stabilization, and protein folding signifies its pivotal role in the infectious process. Furthermore, our endeavor encompassed the categorization of effectors into carbohydrate-active families, illuminating the prominence of Carbohydrate esterases family 12, primarily catalyzing cellulose and xylan degradation. By amalgamating these multifaceted methodologies, our study provides invaluable insights leading to the formulation of more effective ASR molecular strategies, demonstrating its relevance and innovation in the fields of plant pathology and molecular biology.

## 1. INTRODUCTION

Asian soybean rust (ASR), caused by the fungus *Phakopsora pachyrhizi*, is a severe disease with increasing potential due to the expansion of the cultivation area around the world. Yield losses can exceed 90% (Chaloner et al., 2021; Godoy et al., 2016) and the damage is more pronounced in South America where the high costs of chemical treatments generate expenses in the range of billions of dollars annually (Yorinori et al., 2005; Kunjeti et al., 2016). During infection, *P. pachyrhizi* forms specialized structures called haustoria that suck nutrients from the host (Garnica et al., 2014), but also has the role of secreting effector proteins directly into host cells (Petre et al., 2014). These effectors manipulate the soybean immune system, effectively disarming its defenses and allowing the fungus to thrive (Bueno et al., 2022). Thus, effectors are key parts within plant breeding aimed at resistance against phytopathogens and knowing the entire set of them (effectorome) in a given pathosystem provides support for understanding the infection process.

Simplistically, effectors are defined as small molecules secreted into the extracellular space (apoplast) or translocated within the host cell to suppress the immune system and physiology in order to facilitate colonization (Arroyo-Velez et al., 2020; Todd et al., 2022; Todd et al., 2022b). They have specific characteristics such as being smaller than 300 aa, rich in cysteine (>2% cysteine or >4 cys residues), presenting a secretion signal peptide and lacking transmembrane domains (Stergiopoulos; de Wit, 2009; Lo Presti et al, 2015; Carreón-Anguiano et al., 2020). Several biostatistics approaches have already been worked on with a view to identifying potential effectors and their interactions in planta in this pathosystem, the main one being based on transcriptome (Carvalho et al., 2017; Elmore et al., 2020). With the advent of advances in “omics”, especially proteomics, it is possible to obtain the complete proteome of organisms in databases and from there carry out more in-depth studies focused on the set of all effectors of a species (González-Fuente et al., 2020). There is a range of tools that can be used to, from a proteome/secretome, predict candidate effectors, among which SignalP 4.1 (Petersen et al. 2011), WoLFPSORT (Horton et al., 2007), TMHMM 2.0 (Krogh et al., 2001), Secretool (Cortázar et al., 2014) and EffectorP (Sperchneider et al., 2018). However, these types of work in short are limited to showing the number of candidate effectors in different species or observing possible associated functions, but in isolation. None focus on aspects of interaction between these and possible host targets.

Host-Pathogen Protein Interaction (HPPI) studies emerge as a potential for information integration and, together with the effectorome, can provide powerful insights into the plant-pathogen system. This technique, based on Deep Learning, aims to predict possible interactions between host and pathogenic proteins using the concept of Ortholog-based method (or interolog) in which PPI associations (protein-protein interactions) experimentally evaluated in model systems are transferred to non-systematic systems. model or little studied (Matthews et al., 2001). This method has been used by several researchers (Shoemaker; Ar, 2007; Dyer, 2007; Loaiza; Kaundal, 2021; Kaundal et al., 2022) and was first proposed by Yu and collaborators (2004). It assumes that genes that interact in a given pathosystem have orthology across different species, since these genes “coevolve” and are conserved between organisms (Walhout et al. 2000; Yu et al., 2004). Therefore, transferring annotation, within the scope of comparative genomics, between species helps to fill the gap in non-model systems, as well as mitigate the high costs, labor-intensive and time-consuming of in planta experimental analyzes or techniques such as Yeast-Two-Hybrid (Kaundal et al., 2022; Piehler, 2005; Rao et al., 2014).

With this in mind, understanding the precise mechanisms by which *P. pachyrhizi* effectors suppress plant immunity, as well as identifying potential target proteins that are key, is critical to developing effective strategies against ASR. This article aims to integrate effectoromics with HPI networks to uncover the arsenal that this fungus uses to suppress the soybean defense system at the level of PPI interaction. To date, no reports using this type of approach have been found in the scientific literature for this pathosystem.

## 2. MATERIAL AND METHODS

### 2.1. Proteomic retrieval

The respective proteomes of Glycine max (Wm82.a4. v1) and *Phakopsora pachyrhizi* (K8108 v2.0) (Gupta et al., 2023) were retrieved from the Phytozome v. database. 13 (Goodstein et al., 2012) and MycoCosm (Grigoriev et al., 2014), respectively.

### 2.2. Predicted Effectorome of*Phakopsora pachyrhizi*

The fungus proteome was used as input within Secretool (Cortázar et al., 2014) in order to predict the potential proteins secreted by the species. All parameters were used as default. This procedure is necessary to generate the input (secretome) recommended for predicting effectors in the EffectorP tool, since this is not a secretome prediction tool and the maximum number of sequences on the platform is limited to 4000.

To predict candidate effectors from the previously obtained secretome, the EffectorP 3.0 tool (Sperschneider et al., 2021) was used, which is a web server and the most used for this purpose. Sequences in the fasta file were used as input with default parameters. The output generated from the processing was used to generate the HPI network.

### 2.3. Effectorome Subcellular Localization Prediction

Understanding the compartment in which effector proteins are possibly secreted helps to understand potential mechanisms, as well as target molecules that pathogens can manipulate with a view to suppressing the immune system. For this, the machine learning-based tool LOCALIZER (Sperschneider et al., 2017) was used for this prediction. The mechanics of this tool assume that proteins targeted to the chloroplast and mitochondria have a molecular signature (an N-terminal transit peptide), while those targeted to the nucleus have an NLS (Nuclear Localization Signal) (Jarvis, 2008; Holbrook et al., 2016).

The output from the previous procedure was used in Cytoscape to visualize the predicted compartments associated with the effectors.

### 2.4. Network analysis of host-pathogen protein interactions (HPPI) interolog-based

The PPI network between candidate effectors and possible target proteins in soybean was predicted using the web-server for host-pathogen interaction prediction and visualization (PredHPI) (Loaiza; Kaundal, 2021). For this, the “interolog” module, which is based on the model proposed by Yu et al. (2004) and follows the idea already discussed. The soybean proteome was used against candidate effectors of the fungus. All PPI interaction databases available in the tool have been marked. The remaining parameters were used as default. At the end of the process, the output containing the nodes and connections, assigned by orthology to the query sequences, was downloaded so that the underlying network visualization processes, as well as GO enrichments, could be conducted in Cytoscape v. 3.10.1 (Shannon et al., 2003). The enrichment carried out here aimed to attribute possible modules of effectors in association with host proteins. Since the StringDB plugin for annotation in Cytoscape does not have a database for *P. pachyrhizi*, the GO terms were assigned only to the host.

### 4.6 Functional annotation and Carbohydrate-Active enzymes (CAZy) characterization of the ***P. Pachyrhizi*** effectors

In order to obtain greater details at the functional level of individual sequences and thereby gain insights into the way in which effectors act, candidate effectors were subjected to functional annotation using the Blast2GO software (Conesa et al., 2005), which is a universal annotation tool used mainly in organisms lacking functional characterization or non-models. All processes followed the manual pipeline for this type of analysis which includes, among others, Blast of sequences, mapping, annotations and presentation of Charts. In the Blast option, we chose to use the NCBI web service. In the alignment settings, the analysis was restricted to the fungi taxon, as well as parameters were defined for the number of Blast Hits equal to 20, E-value (1.0E-05), word Size equal to 7 (amino acids) and also the filter for low complexity regions was checked. All other parameters were kept as default.

Aiming to identify, within the candidate effectors, those with carbo-active enzymatic function, we used the Carbohydrate-Active enZYmes (CAZy) database (Drula et al., 2022) through the dbCAN3 (Automated Carbohydrate-active enzyme ANnotation) software (Zheng et al., 2023). This tool uses both the CAZy database and the HMMER-based method to classify carbo-active enzymes into families, as well as provide possible substrates on which they act. Simply put, in the Annotate module, the amino acid sequences of the effectors were used as query and the tools HMMER: dbCAN, DIAMOND:CAZy and HMMER: dbCAN-sub were marked. Only those effectors predicted by at least two of these three algorithms were retained in the subsequent analysis for greater reliability.

## 3. RESULTS & DISCUSSION

### 3.1. Effectorome prediction and subcellular localization

The effectorome of *P. pachyrhizi* (K8108), studied here, was composed of 185 candidate proteins predicted as effectors. In general, the number of effectors varies between different pathogens. This variation is related to both lifestyle (biotrophic, hemibiotrophic or necrotrophic) and occurs even within the species. A study comparing 40 phytopathogens showed, for example, that the size of the effectorome varied between 708 (Auricularia subglabra) and 26 (Saccharomyces cerevisiae) effectors, with an average of 400 for hemibiotrophs, 200 for necrotrophs and 300 for biotrophs (Carreón-Anguiano et al., 2020). The study also suggests that the number is linked to the complexity of interaction between plants and hosts for these classes, as there is a need for a greater repertoire and specialization for the invasion and colonization of tissues while still keeping the organism alive.

Our result is in agreement to those obtained using transcriptome. Link et al. (2014), using haustorium transcriptome, identified 156 candidate effectors in the *P. pachyrhizi* isolate Thai-1. Exemplifying this intraspecific divergence, isolate GA-05 presented only 36 (Kunjeti et al., 2016). This suggests that there may be a positive selective pressure for these effectors that tend to evolve with the niche and that possibly, in the sense of coevolution, have different sets of effectors used as strategies to suppress host defense.

Another important step in a study of this type is knowing the subcellular location of these effectors. Effector proteins can be secreted into extracellular spaces (apoplast) or within the cytoplasm (intracellular). In the latter case, the objective is to attack proteins within the cytoplasm and cellular organelles (Figueroa; Ortiz; Henningsen, 2021), which, once inside the spaces, are directed to the various compartments such as the nucleus, plasma membrane, chloroplasts, mitochondria (Lindeberg et al., 2012; Deslandes; Therefore, knowing the localization in plant cells helps to understand how the infection program is engineered by the microbe (Sperschneider et al., 2017). In this study, only predictions with associated probabilities above 70% (0.7) were considered. It is worth noting that EffectorP does not provide probability for nuclear predictions, therefore, although connections in red may indicate confidence of 0.90-0.95 (following the colors in the legend), they should be ignored (only for nuclear location predictions). Following this parameter, figure 1 shows the result in network. The number of proteins localized to the chloroplast with transit peptide (cTP) was 19, with a total of 33 (17.8%) of the effectors being present in this compartment; mitochondrial ones with transit peptide (mTP) were 12 (6.5%). Effectors with exclusive nuclear localization and NLS signal represented 16 (8.6%) and with both NLS and TP (transitpeptide) accounted for 10 (5.4%) of the 185 effectors submitted for analysis.

**Figure 1.**
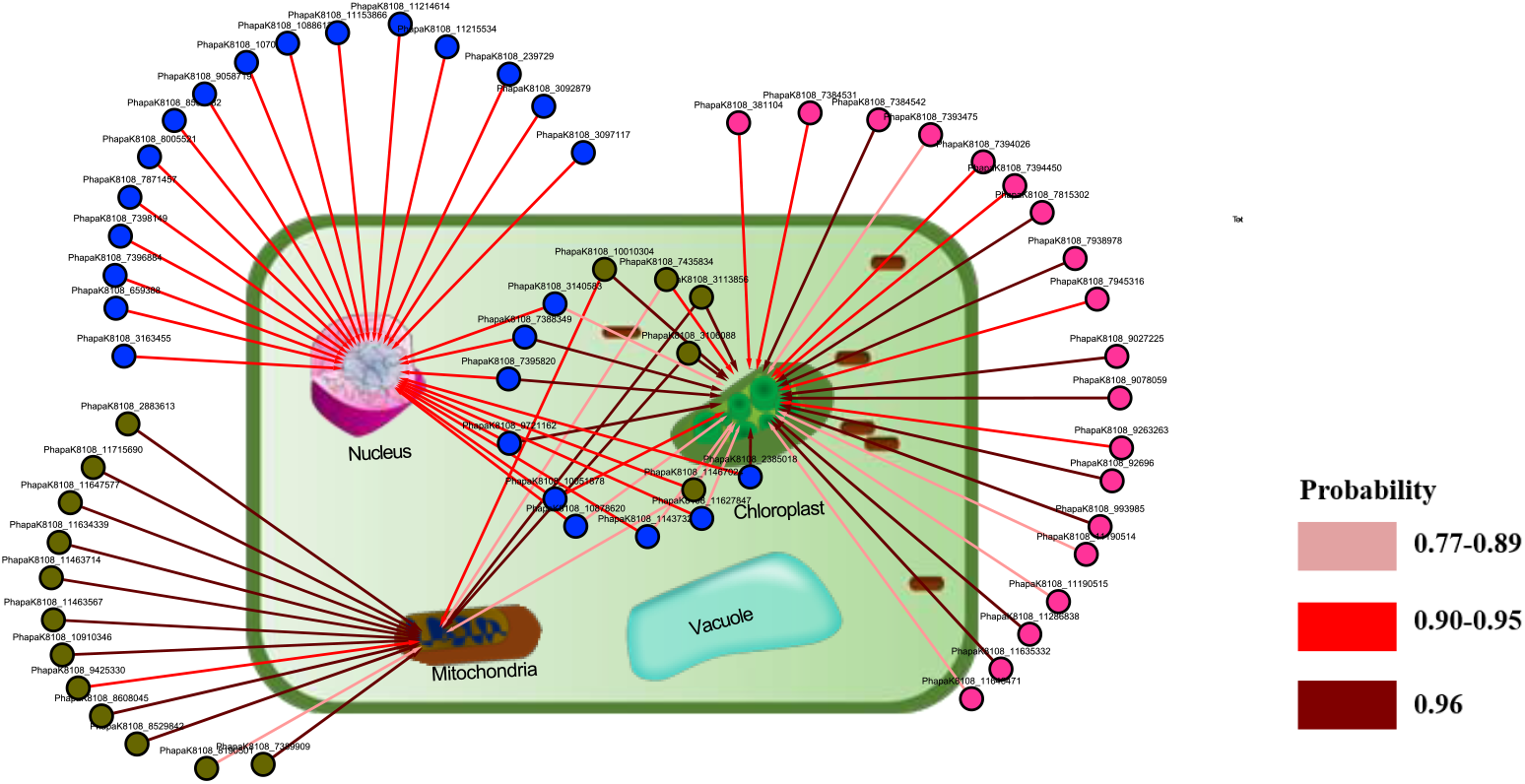
Subcellular localization prediction network. Relational illustration between the Nucleus, Mitochondria and Chloroplast compartments and their corresponding predicted effectors (nodes colored according to the address). The nodes outside the cell represent those with NLS (Nucler localization signal), cTP (chloroplast transit peptide), mTP (mitochondria transit peptide), respectively for the colors Blue, Pink and dark green. The nodes within the cell have more than one location indicated by arrows. The thickness of the connections, as well as the intensity of red, proportionally indicates the probability indicated in the legend.

Only four effectors (PhapaK8108_3106088, PhapaK8108_3113856, PhapaK8108_7435834, and PhapaK8108_10010304) had cTP and mTP at the same time; only the effector PhapaK8108_11467024 presented TP (for chloroplast and mitochondria) and NLS, therefore the only one addressed to the three compartments. As found, the effectorome (composed of 185 effector proteins) presented 71 predicted proteins in at least two cellular compartments. Indicating a large repertoire and potential targets within the host.

There are major questions about the ways in which fungal/oomycete effectors cross the host membrane. Results obtained with *Melampsora larici-populina* suggest that a possible mechanism occurs by mimicking host TPs to direct these proteins to organelles (Petre et al., 2016), that is, using the organism’s molecular signatures to usurp the cellular machinery to its benefit. Fact is, once the effector proteins (EPs) are inside, they use these host molecular signatures to gain access to the organelles. To illustrate, some EPs have, in their sequences, domains that mimic the host’s NLS, consequently they are addressed to the nucleus through nuclear pores (Schornack et al., 2010; Wirthmueller et al., 2015) as if they were regular molecules of the host system. In this study, 26 effectors (nodes in blue, plus the node representing PhapaK8108_11467024) were predicted to be targeted to the nucleus. Both the nucleus and chromatin are preferred targets for the pathogenic arsenal with a view to manipulating the transcriptional profile and important defense response routes such as the hormonal signaling pathway and MPAK (Baxt et al., 2013). In the context of chromatin, they manipulate the epigenome in order to prevent the host surveillance system from detecting the invader (Locatelli et al., 2023; Monerri; Kim, 2014), in this case modulating the expression of important genes in the chromatin pathways. response as plant hormones. Therefore, it is reasonable to think about the importance of EPs directed to the nucleus within the infection process and due to the number of effectors identified here targeting the nucleus, we see the aggressiveness of this fungus towards the host.

In addition to the action at the epigenetic level, the transcriptional level is also greatly affected. Numerous effectors of phytopathogens such as *Pseudomonas syringae*, *Hyaloperonospora arabidopsidis* (Hpa), and *Phytophthora infestans* have been reported to interfere with host transcriptional reprogramming by interacting with NAC-type transcription factors (TFs) (Block et al., 2014). These TFs are key players within the plant immune system for their role in the SA (salicylic acid) signaling pathway, ROS modulation, endoplasmic reticulum (ER) stress, as well as PCD (Programmed cell death) (Reis et al., 2016; Yang et al., 2014). This shows one of the important strategies for addressing EPs from microbes to the nucleus and its role in host transcriptional deregulation, which is in no way disconnected from epigenomics. The defense system has several layers that communicate and sometimes even overlap, making it even more difficult to implement a direct strategy when aiming to develop resistant cultivars. It is worth highlighting that the identification of effectors acting on a process only concerns systems already studied experimentally. The actual and observed quantity may be many orders of magnitude greater. This is also true on the host side when observing some interacting genes or targets of pathogens.

### 3.2. Functional annotation and Carbohydrate-Active Enzymes (CAZyme) characterization of the***P. pachyrhizi***effectors

The analysis of effectors with carbo-active activity additionally identified (fig. 2) the family and substrates on which each one acts.

**Figure 2.**
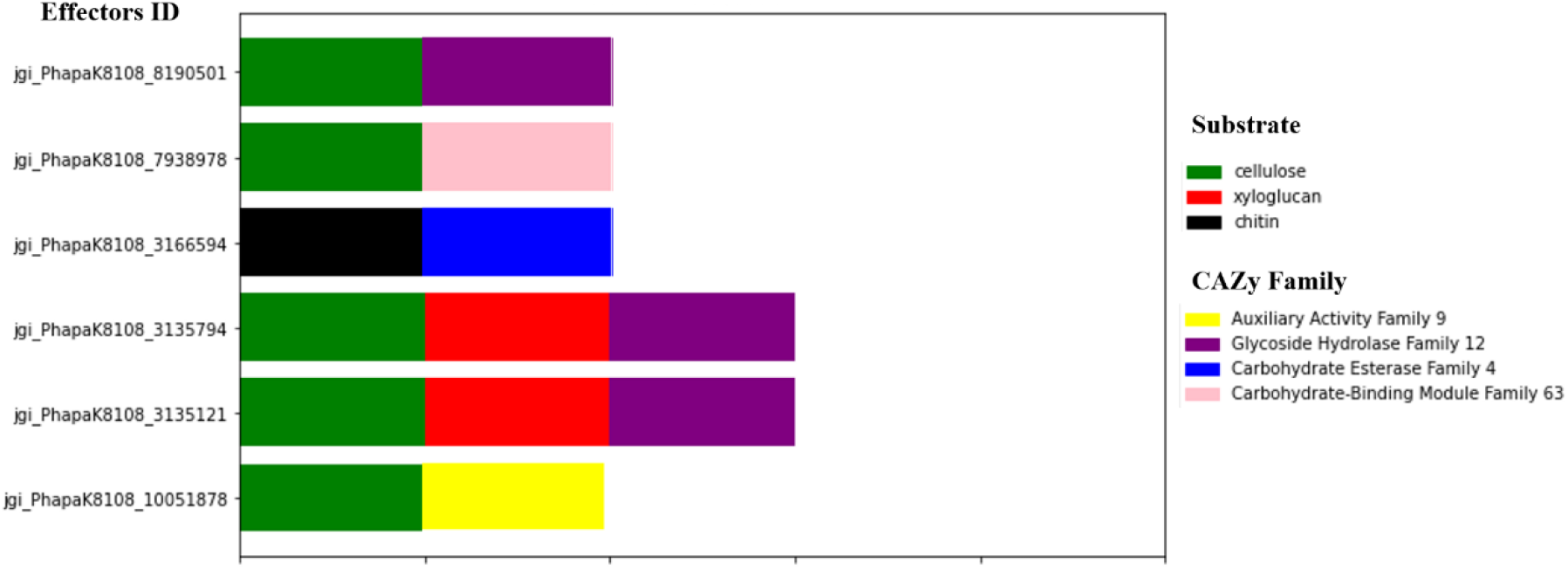
Identification of Carbohydrate-active enzymes (CAZymes), families and substrates from the effectorome. On the X axis, the families and substrates related to each of the IDs of the genes that encode the enzymes are positioned in different colors.

In total, 13 enzymes with carbohydrate activity were identified, six of which were confirmed in both tools (Blast2GO and dbCAN3) and only these are presented in the figure (the additional ones can be seen in the supplementary material: tab 1), since the enzymes identified by Blast2GO do not show the acting substrate. Carbohydrate-active enzymes (CAZymes) are extremely important within the infection program carried out by filamentous phytopathogens. This is because they participate in several processes that culminate in the invasion, colonization and finally dispersion of the pathogen in a positive interaction with the host (Vries; Vries, 2020). Rusts need to access the internal part of the host cell, which in turn has cell walls protected by compounds such as cellulose and pectins that need to be degraded. Therefore, these fungi secrete enzymes such as Glycoside Hydrolase (GH) and esterases that perform this role and are considered cell wall degrading enzymes (CWDEs) (Heller, 2020; Mafa et al., 2023). Our results show that two candidate effectors (jgi_PhapaK8108_3135794 and jgi_PhapaK8108_3135121) are proteins from the GH family 12 and have cellulose and xyloglycan as substrates. The latter is the structural polysaccharide of great importance in the primary cell wall of higher plants and has the function, among others, of providing rigidity (Hayashi, 1989; Eckardt et al., 2006; Zheng et al., 2018), which is why they become interesting targets for the pathogen. According to CAZy, members of this family have Xyloglucan-specific endo-β-1,4-glucanase activity, being capable of degrading glycans and cellulose (Grishutin et al., 2004; Lombard et al., 2014). Therefore, these two EPs are good candidates to play a relevant role in the initial infection process by suppressing PTI (PAMP-triggered immunity).

Five of the six candidate effectors presented in the figure have common substrates, that is, cellulose. However, it is interesting to note that despite this characteristic, the families to which they belong are distinct. In addition to the GH family, they comprised Auxiliary Activity family 9 (AA9) and Carbohydrate-binding module Family 63 (CBM63). This suggests that different actors participate in the deconstruction of these carbo-compounds. And in fact, the process to degrade cellulose is much more complex and takes place in stages. There is a need for the action of four different enzymes acting in the primary phase and for complete degradation, auxiliary enzymes come into play that hydrolyze the side chains of the compound (Claes et al., 2020; Minty and Lin, 2015). The AA9 family identified in this study is considered one of the enzymes with an auxiliary role, not acting directly on carbohydrates, but jointly solubilizing lignin fragments that are invariably found together with carbohydrates such as cellulose (Levasseur et al., 2013). For example, jgi_PhapaK8108_10051878 has Lytic cellulose monooxygenase (LPMOs) activity, oxidizing recalcitrant carbohydrates such as cellulose. In the process, it breaks down polysaccharides such as cellulose, creating fission points that culminate in the disintegration of the fibrillar structure (Villares et al., 2017). Taking into account that the cell wall is the first defense barrier that the host has against microbes, the importance of these enzymes in suppressing the basal system of plants at the PTI level is paramount. LPMOs play an even more relevant role because cellulose, the main constituent of the primary wall, and target for degradation, has a crystalline structure that can only be destabilized with the action of oxidative enzymes of this type. Together, both LPMO and CAZymes work in coordination where the former acts to destabilize and the latter to depolymerize (Jagadeeswaran; Veale; Mort, 2020). It is noted that the level of specialization of the pathogen to avoid the first defense barrier is very high and several gene products act together.

The other family that can be part of this auxiliary process is the Carbohydrate-Binding Module Family 63 (CBM63). Just like AA, CBMs take part in CAZymes complexes with a view to increasing the degradation potential of the target (Abbott; van Bueren, 2014), by increasing thermosensitivity (Krska; Larsbrink, 2020) or even promoting increasing the concentration of enzymes around the substrate and consequently the reactive rate (Hervé et al., 2010). In addition to this, its main function is to recognize and bind carbohydrates (Boraston et al., 2004; Duan et al., 2016), and like LPMO, it also destabilizes the structure on recalcitrant substrates, increasing the action of CAZymes (Wang; Zhang; Gao, 2008). CBM63 is a type of expansins that binds cellulose (CAZYpedia database) with high affinity to the cell wall (Georgelis et al., 2011). The results of this latest article suggest that the two structural domains (D1 and D2) of this enzyme are not only important in anchoring to the surface of the cell wall, but also play a role in loosening the wall, which would be a beneficial strategy for the pathogen. Something quite interesting is that this type of proteins, according to the CAZYpedia database, was incorporated into the genome of phytopathogens via horizontal transfer with plants. This may imply that they use, throughout evolution, elements of the host itself to their advantage in order to escape the host surveillance system.

The effector jgi_PhapaK8108_3166594 was the only one to have chitin as a substrate and belongs to family 4 of Carbohydrate Esterase (CEs). The main activities related to this member are Chitin oligosaccharide deacetylase and Chitin deacetylase (these and other activities can be seen in the supplementary material: tab.1_supplementary). Therefore, they act mainly as chitinases. Although, in fact, plants do not have chitin, in pathosystems that have mammals as hosts, the pathogen used an alternative substrate structurally similar to chitins to establish the interaction (Chaudhuri et al., 2010). It can use, for example, a GlcNAC residue (β-1,4-N-acetyl-D-glucosamine) from glycoproteins as a substrate for chitinase (Lin et al., 2020). In general, these secreted proteins play a primary role in cell adhesion during the early stages of pathogenesis. Furthermore, they are important factors for virulence (Jeong et al., 2023). Considering that the adhesion of spores to the host surface is an imperative step in the infection process, it can be argued that it becomes a key factor of positive interaction and can define the infection process (Asadi et al., 2019) and which can be the subject of further study, for example, loss-of-function to confirm.

In addition to the aforementioned families, three other effects (JGI_PHAPAK8108_916471, JGI_PHAPAK8108_381104 and JGI_PHAPAK8108_11153866) were identified as belonging to family 3 of Polysaccharide Lyase (PL3) that are used to use cell walls (Cnossen-Fassoni et al., 2013), mainly pectin (Wan et al., 2021). Chemically speaking, its action occurs by randomly cleaving α-1,4-polygalacturonic acid, affecting the plant cell wall. Deletion of genes encoding the enzyme resulted in lower virulence (Collmer, 1986; Yakoby et al., 2001; Ben-Daniel et al., 2011). A similar gene inactivation approach was conducted with *Colletotrichum lindemuthianum* and as a result, less aggressiveness in anthracnose symptoms was observed (Cnossen-Fassoni et al., 2013). On the other hand, PL3 are considered both virulence factors and elicitors of PTI. For example, a member of this family was studied in *Fusarium sacchari* showing that, in addition to inducing cell death, the protein’s signal peptide functioned as a molecular signature for the host to trigger PTI (Wang et al., 2023). This suggests the dual role of this effector within this pathosystem. The same was reported in the *Verticillium dahliae*-Tobacco pathosystem (Yang et al., 2018).

In addition to the EPs identified and characterized in terms of family, substrate and main activity, three others were exclusively obtained by functional annotation via Blast2GO (supplementary material:Blast2GO_results). These are candidate effectors jgi_PhapaK8108_3113856, jgi_PhapaK8108_7408047 and jgi_PhapaK8108_3062326. Its activities were associated, respectively, with superoxide dismutase, cyclophilin-like domain and Aldehyde/histidinol dehydrogenase. The annotation attributed the biological process (BP) to jgi_PhapaK8108_3113856 as removal of superoxide radicals; protein folding for jgi_PhapaK8108_7408047 and jgi_PhapaK8108_3062326 exclusively the molecular function of oxidoreductase. Due to the relevant biological aspect, the effector jgi_PhapaK8108_7408047 will be discussed in the HPPI section, as it has proven to be a key component in the network.

An addendum is needed regarding EPs involved in superoxide radical (SOD) scavenging (jgi_PhapaK8108_7408047). This enzyme is part of the cellular apparatus used to protect the system against oxidative stress present in the form of superoxide radicals (Oztürk-Urek; Tarhan, 2001). Thus, the harmful potential of these agents of exogenous origin is eliminated by the presence of SOD, which are antioxidants (Bannister; Rotilio, 1987). Taking into account that one of the first lines of defense is oxidative stress (ROS), which destabilizes essential components of the pathogen such as proteins and the cell wall (Haas; Goebel, 2002; Hasset; Cohen, 1989), it would be reasonable to think that SOD would act by removing potential radicals produced by the host. And in fact, there are studies that show SOD is required for the virulence of *Pseudomonas aeruginosa* on *Bombyx mori* (Liyama et al., 2007), therefore, it represents the first line of defense against the stress caused by superoxide, converting it to peroxide of hydrogen and oxygen, thus protecting cells from the toxic effects of superoxide (Iiyama et al., 2007). Another potential detoxifying agent present in the arsenal of *P. pachyrhizi* is the effector jgi_PhapaK8108_3062326, which we identified as having Aldehyde/histidinol dehydrogenase activity with a molecular oxidation reductase function. Aldehyde dehydrogenase, as well as SOD, are collectors of oxidative species present in the medium (Felczak; TerAvest, 2022). Gene deletion in *Magnaporthe oryzae* impacted pathogenicity (Abdul et al., 2018). Therefore, it appears that *P. pachyrhizi* makes use of effectors with roles that sometimes overlap functions in order to enhance its colonization.

Taken together, the results thus far highlight the complexity of the interactions between *P. pachyrhizi* and G. max. The identification and characterization of carbo-active enzymes reveal a wide variety of mechanisms used by pathogens to overcome the plants’ first (structural) defense barrier and facilitate infection. Furthermore, the diversity of candidate effectors and their activities suggests a highly organized strategy adopted by phytopathogens, ranging from cell wall degradation to modulation of the plant’s immune response (transcriptional/epigenetic modulation). These insights can provide a crucial basis for understanding the infection process, as well as opening new perspectives for understanding the coevolution between plant pathogens and their hosts.

### 3.3. Effectorome network analysis of Host-pathogen Protein Interactions (HPPI) interolog-based

Apart from the potential effectors identified with carbo-activity, a more global view of protein-protein interactions (PPI) between *P. pachyrhizi* effectors and the entire G. max proteome would provide powerful information on possible proteins targeted by the pathogen with a view to cause disease. With this in mind, and within the scope of interolog-based prediction, the host-pathogen interaction network (HPPI) seen below was obtained (Fig. 3).

**Figure 3.**
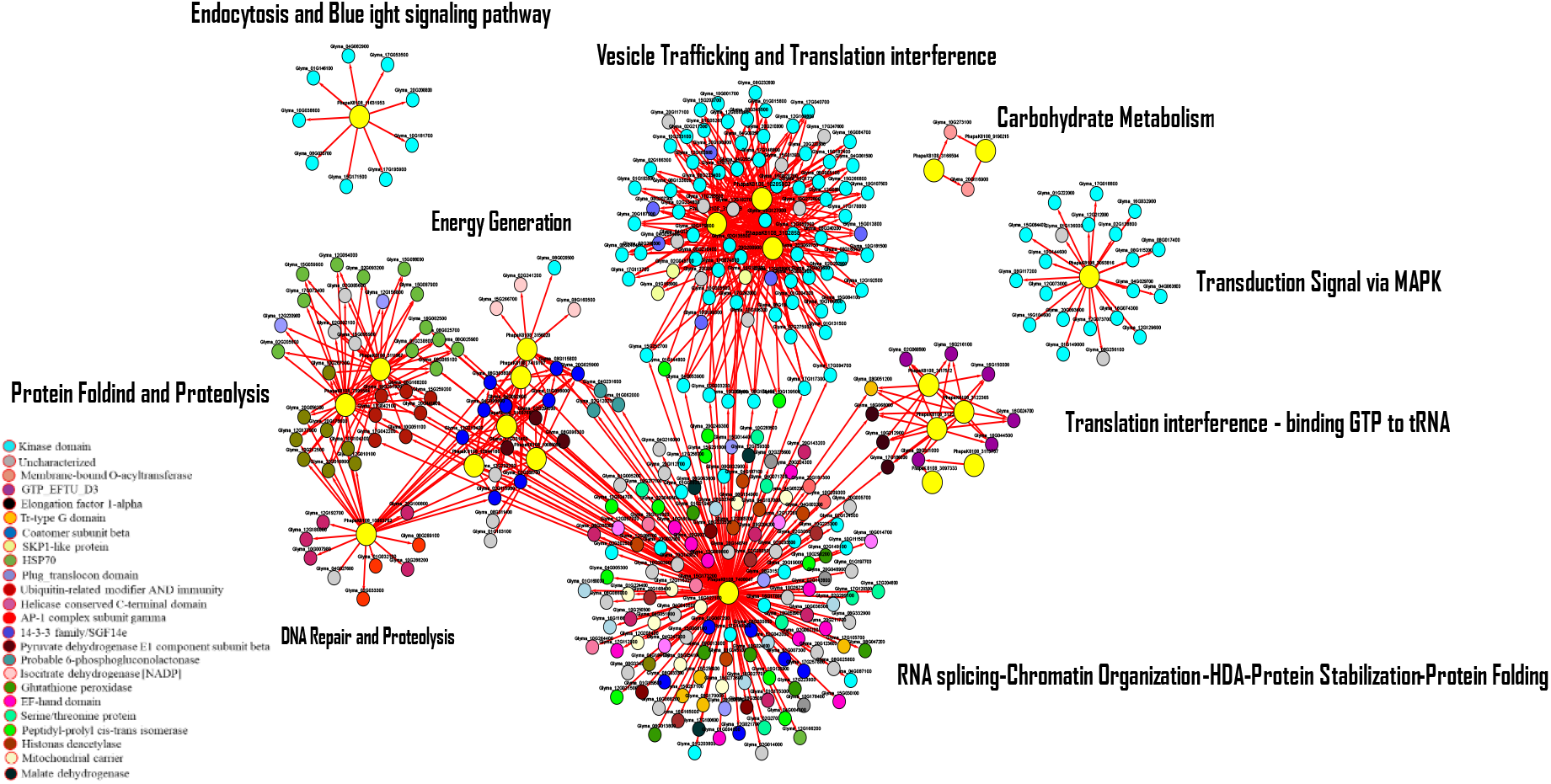
Interolog-based HPPI network. The larger nodes (yellow) are the phytopathogenic effectors; Smaller nodes in different colors represent different classes of target proteins in G. max. The captions next to each cluster indicate the associated modules. The connections have arrows that point toward target proteins in the host. The smaller (separate) subnetworks indicate proteins with more specific interactions and are not directly related to the interactions of the larger network. Predominant protein class/biological process is labeled (bottom left). The colors respectively represent the protein classes to which the group of nodes belongs. Proteins considered Unknown are gray in color. Labels: GTP_EFTU_D3 for (elongation factor Tu (EF-Tu) GTP binding domain: Pfam: PF00009); Tr-type G domain (Translational (tr)-type GTP-binding domain); SKP1-like protein (S-phase kinase-associated protein1); HSP90 (Heat shock protein 70)

The PredHPI tool predicted 922 unique PPIs among the 185 candidate effectors from the fungus and the soybean proteome. This means that this number was the total number of interactions found by orthology (interolog) within the experimentally evaluated and curated databases in which the interactions of a model are used to predict the interaction in new systems (Nourani; Khunjush; Durmu, 2015 ;Matthews, 2001). Several computational methods like this have already been used for this purpose, differing only in the approach, for example, using amino acid sequence homology and domains (Pellegrin et al., 1999; Yu et al., 2004; Wu et al., 2006 ; Garcia-Garcia et al., 2012; Therefore, it is a powerful approach to generate PPI networks through intraspecific transferability.

In total, 344 orthologous proteins from both organisms were identified within the databases, 324 belonging to the host and 21 to the pathogen. For the pathogen, the largest class was Lysophospholipases (LysoPLs) with five effectors (PhapaK8108_3097333, PhapaK8108_3113757, PhapaK8108_3117512, PhapaK8108_312271, PhapaK8108_3122365), followed by subtilisin protease (PhapaK810 8_10285803, PhapaK8108_3102858, PhapaK8108_3102669) and Glycoside Hydrolase (PhapaK8108_7419107, PhapaK8108_3109768, PhapaK8108_10944186) with three effectors each. In addition to these, we identified an uncharacterized protein (PhapaK8108_11631953), an Acid protease (PhapaK8108_10863762), Aorsin (PhapaK8108_10085059), Carbohydrate esterase (PhapaK8108_3166594), Carboxypeptidase (PhapaK8108_3111967, paK8108_7395564), Cyclophilin-like (PhapaK8108_7408047), Glucanase (PhapaK8108_3093816), Peptidoglycan deacetylase (PhapaK8108_9190215) and Phosphatidylethanolamine-binding protein PEBP (PhapaK8108_3156020). More details (supplementary material)

Regarding the number of proteins they interacted with, the candidate effector PhapaK8108_7408047 (Cyclophilin-like) was the one that targeted the largest number of proteins (171), both PhapaK8108_3097333 and PhapaK8108_3113757 (Lysophospholipase) had only 1 and were the smallest. PhapaK8108_3102669, PhapaK8108_3102858 with 87 and PhapaK8108_10285803 with 83 were the second and third highest, respectively.

Taking into account the diversity of target proteins and possible biological processes in which this effector takes part and influences, PhapaK8108_7408047 not only had unique interactions within the “mix” module containing several processes (RNA splicing, Chromatin Organization, Histone Deacetylase, Protein Stabilization, Protein Folding), which included 130 proteins, as well as interaction with four other modules (DNA Repair and Proteolysis; Protein folding and Proteolysis; Energy Generation; Tranlation interference; Vesicle trafficking). Below (fig. 4) this effector is highlighted. The results indicate that it may be an important effector within the pathogenic arsenal, as it targets different processes as well as a broad spectrum of proteins.

**Figure 4.**
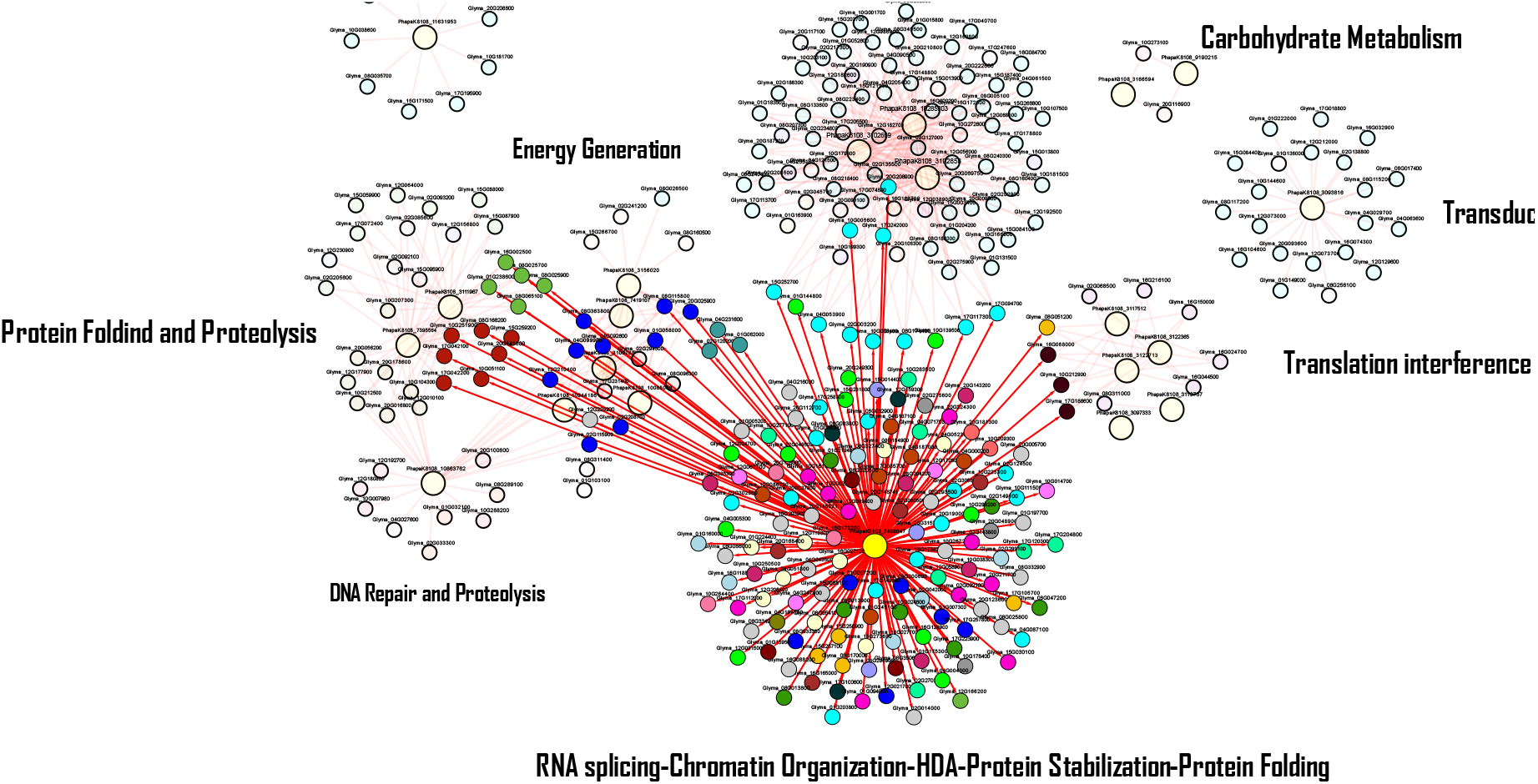
Effector PhapaK8108_7408047 highlighted (larger node in yellow) interacting with the largest/diverse module (RNA splicing-Chromatin Organization-HDA-Protein Stabilization-Protein Folding). Colored nodes represent target proteins in different modules. Open nodes indicate proteins that do not directly interact with this effector.

For example, within the interference module in energy production processes, interaction was observed with host proteins of the 14-3-3/SGF14f type, which act as phosphopeptide-binding proteins and target proteins related to transduction genes. of signals (Darling et al., 2005; Gokirmak et al., 2010).

The involvement of this target protein in the regulation of the transcriptional network has already been reported. For example, they can interact with transcription factors such as Basic leucine zipper and tissue-specific regulatory factor to possibly regulate the abscisic acid (ABA) signaling pathway (Schultz et al., 1998). In addition to the interaction with the transcriptional apparatus, our results corroborate the role of PhapaK8108_7408047 at the post-translational level. Proof of this is that within protein folding and proteolysis, the interaction was carried out through seven Ubiquitin-like proteins and five HSP70 proteins (Heat Shock 70 kDA Proteins).

Manipulation of the ubiquitination system, which is a post-translational modification that regulates almost all signaling networks within the cell and programmed cell death, is one of the targets of microbial effectors with a view to evading host responses mediated by Ubiquitin or even use it to your advantage (Mukherjee; Dikic et al., 2022). This process leads to the chemical marking of the protein that will be destined for degradation (Ravid; Hochstrasser, 2008), in addition, the assembly, stabilization, maturation of proteins and their complexes are related functions (Wang et al., 2004). This suggests that the pathogen could influence the degradation of proteins important for host defense within the signaling cascades or even prevent the recognition of phytopathogenic molecules. In the latter case, it would act by influencing the processing and folding of NB-LRR (nucleotide-binding domain-leucine-rich repeat) type proteins, which are very specific receptors that recognize phytopathogenic effector proteins (Shirasu, 2009). Following this reasoning, the change in conformation of these receptors would cause the effectors to go unnoticed by the host surveillance system. Related to targeting the host protein for degradation or relocalization, studies have already demonstrated that the HopI1 effector from Pseudomonas syringae can bind to the host’s HSP70 and direct it to the chloroplasts. It was later confirmed that hosts lacking these proteins in the cytosol were more susceptible (Jelenska et al., 2007, 2010). Thus, the importance of this effector in manipulating key recognition processes becomes clear and it appears that, since it is an isomerase, it is done by changing the shape of these target proteins, which then interferes with the natural processes of the cellular machinery.

In addition to these modules under which the PhapaK8108_7408047 effector acted, the other two were Translation interference and Vesicle Trafficking and Translation interference. In the first, the PPI interaction was made with host proteins EF1 (Elongation Factor 1) and Tr-type G domain; in the latter they were Non-specific serine/threonine protein kinase. These two components are involved in the initial processes of protein translation. EF1 ensures proper translation of the information contained in the mRNA, while the Translational (tr)-type GTP-binding domain (Tr-type G) is a release factor of the nascent polypeptide chain from the ribosome. It is interesting to imagine the molecular aspects under which this interference in the translation elongation and termination process takes place, and more than that, which protein(s) would be the target(s), would be an object of study interesting future. In this same module, the five representatives of LysoPLs are present (fig. 3). LysoPLs are types of esterases that hydrolyze lysophospholipids and participate in processes such as lipid catabolism, membrane stabilization and immunity. In many pathosystems, they are considered key elements of virulence, taking part mainly in the host infection process (Hervé et al., 2023). Since it was not possible to establish a direct relationship between these types of effectors and the predicted host proteins, it can be considered that predHPI has not, in this case, been efficient in this attribution or there are still no studies that relate lysophospholipid hydrolysis and translation release and elongation factors. Therefore, there is a lack of information in the literature.

An important target of pathogens is the vesicle trafficking system. This is because a large part of the communication between the different internal compartments of the cell, between cells and with the surrounding environment is carried out through this route. This system, through endocytosis, has the ability to internalize vesicles from sources external to its membrane and this process is what allows the interaction between microbes and plants. Once the effectors are released into the environment, this system absorbs them and the pathogen is able to modulate the host’s immune system (Abubakar et al., 2023). The HPI network shown here (fig. 5), along with additional enrichment processes, is the module mostly involved in this aspect.

**Figure 5.**
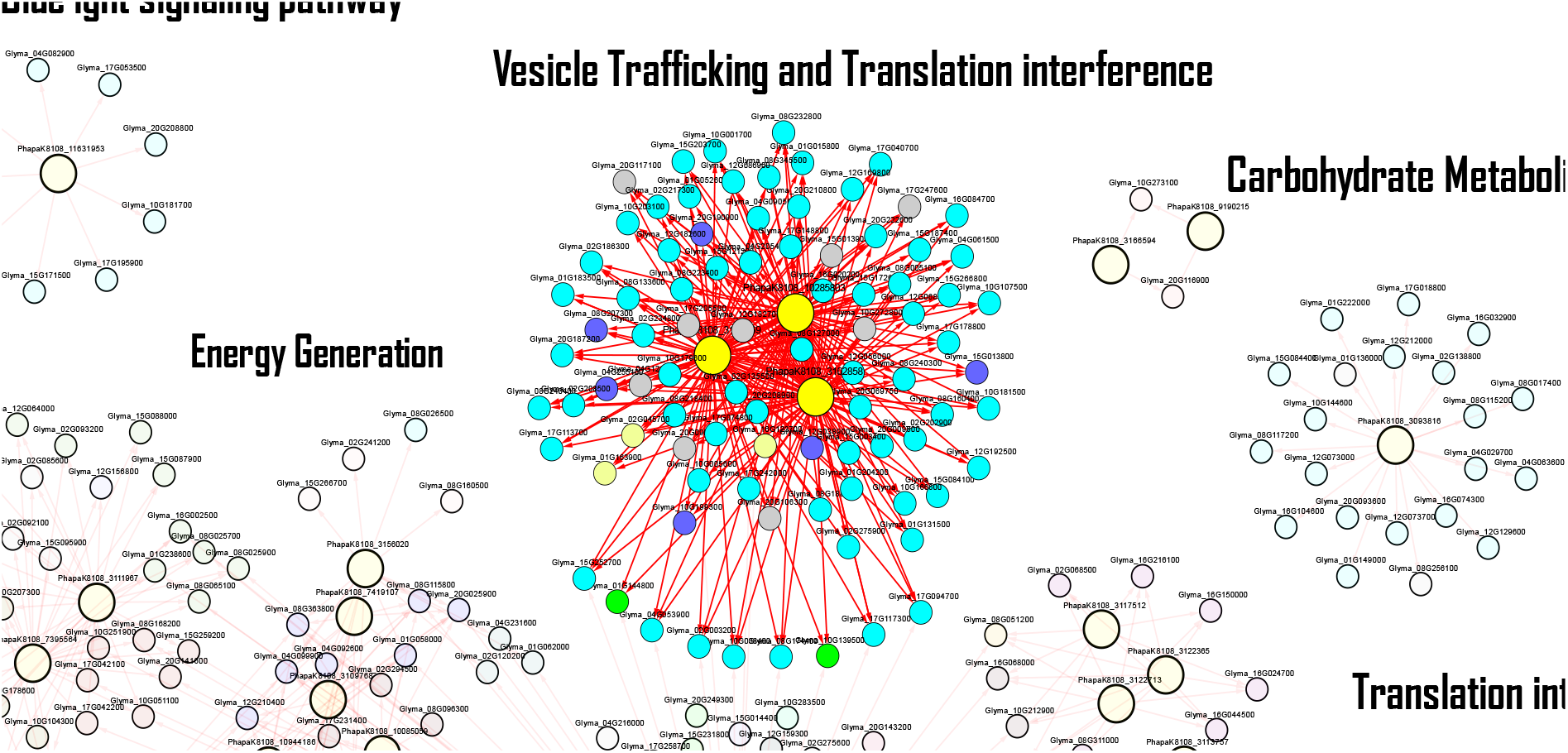
Vesicle Trafficking and Translation interference module. Nodes in yellow show pathogen effectors; the rest are target proteins in the host. Only the module in question was highlighted.

In it, three candidate effectors (PhapaK8108_10285803, PhapaK8108_3102669, PhapaK8108_3102858) had interactions with important actors in this system such as SAC domain-containing protein, which is a membrane protein primarily in the endoplasmic reticulum (Whitters et al., 1993; Foti et al., 2001), responsible for the organization of the cytoskeleton, functions within the Golgi apparatus and vacuole morphology (Zhong; Ye, 2003). This domain has phosphoinositide phosphatase activity, which in turn has metabolism regulated by kinases and phosphatases that phosphorylate and dephosphorylate, respectively (Zhong; Ye, 2003). This explains, in part, the quantity of proteins with kinase domains found in this module (blue nodes). There were 26 types of Non-specific serine/threonine protein kinase (26) and Protein kinase domain-containing protein (31). Members of the serine/threonine protein kinase are essential for triggering the cascade of kinases used to perceive changes in the environment (Hardie; Carling, 1997), and so it is reasonable to think that the destabilization of this super family is beneficial to phytopathogenic colonization, which justifies the total found on the HPI network. Of the 324 proteins attributed to the host, 112 belong to this category. It appears that the pathogen not only targets essential proteins in the membrane structure that enable vesicular trafficking, but also disrupts the kinase-mediated signaling pathway.

Other proteins linked to vesicular trafficking were eukaryotes WD40-repeat (WDR) which has functions similar to the previous one, in addition to involvement in the production of cell wall polysaccharide, assembly of large transmembrane complexes, having the status of supramolecular “hubs” (Guerriero; Hausman; Ezcurra, 2015). In addition, an Inositol polyphosphate phosphatase protein (IPPc) and six Coatomer (purple nodes) closed the list of possible targets related to interference in vesicle trafficking made by the mentioned effectors. IPPc, in addition to being involved in this process, also regulates serine-threonine phosphatases, transcription, mRNA transport, regulation of the cytoskeleton and even cell death (Barker; Berggren, 2013; Livermore et al., 2016; Trung et al., 2022; Shears, 2018; Insall; Weiner, 2001; Piccolo et al., 2004), therefore, several aspects related to the plant’s immune response regulation. In relation to Coatomer, which are complexes of proteins involved in intramembrane transport between ER and Golgi, it is composed of seven units that include, among others, the alpha and beta units (Sánchez-Simarro et al., 2020). In this work, we identified four alphas and 2 betas. The other proteins participating in this and other modules not covered in the scope of this study will be made available in the supplementary table. The module with the highest number of effectors (PhapaK8108_3109768, PhapaK8108_7419107, PhapaK8108_10944186, PhapaK8108_10085059, PhapaK8108_3156020) was represented by metabolism linked to the interference of energy (Energy generation) (fig. 6). The first three are Glycoside hydrolase, followed by Aorsin and Phosphatidylethanolamine-binding protein (PEBP). PhapaK8108_10944186 (Glycoside Hydrolase) interacts only with SGF14f proteins (navy blue nodes), whereas Aorsin, in addition to those mentioned, also interacts with Pyruvate dehydrogenase E1(brown), Transket_pyr domain-containing protein and Glutamyl-tRNA(Gln) amidotransferase subunit A (gray).

**Figure 6.**
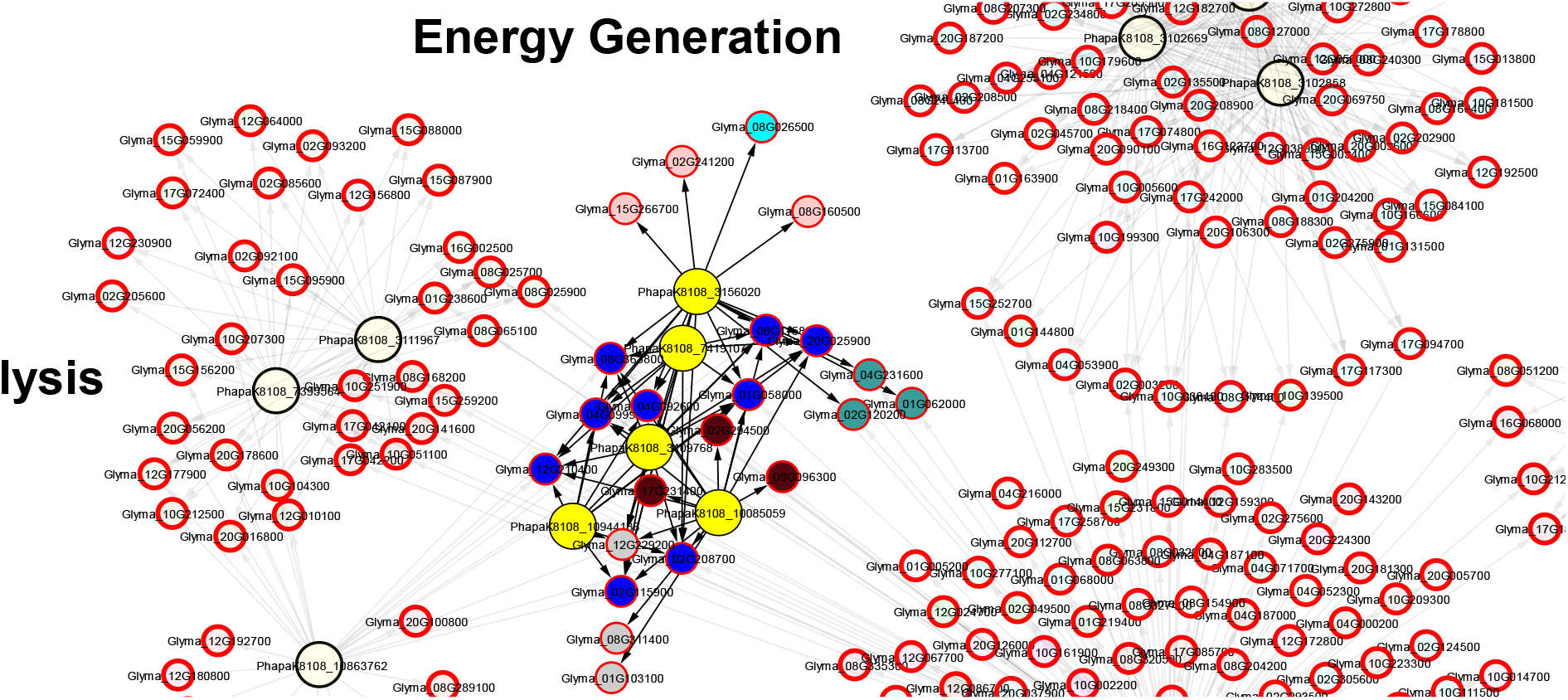
Module related to energy metabolism pathway. In yellow are the identified effectors

It is interesting to note that SGF14f also makes contact with the effector of the DNA repair module, which is an Acid protease. Considering the aspects discussed about the importance of GHs in the infection process, we will focus on the other points. Aorsin is a type of serine proteinase with specific substrate and specific pH to act (Lee et al., 2003). There is very little, or even no, information related to aorsin as a virulence factor. Furthermore, these target proteins GF14f/14-3-3) showed a greater interaction diversity. Theyinteracted with seven effectors from three different modules (fig.7).

**Figure 7.**
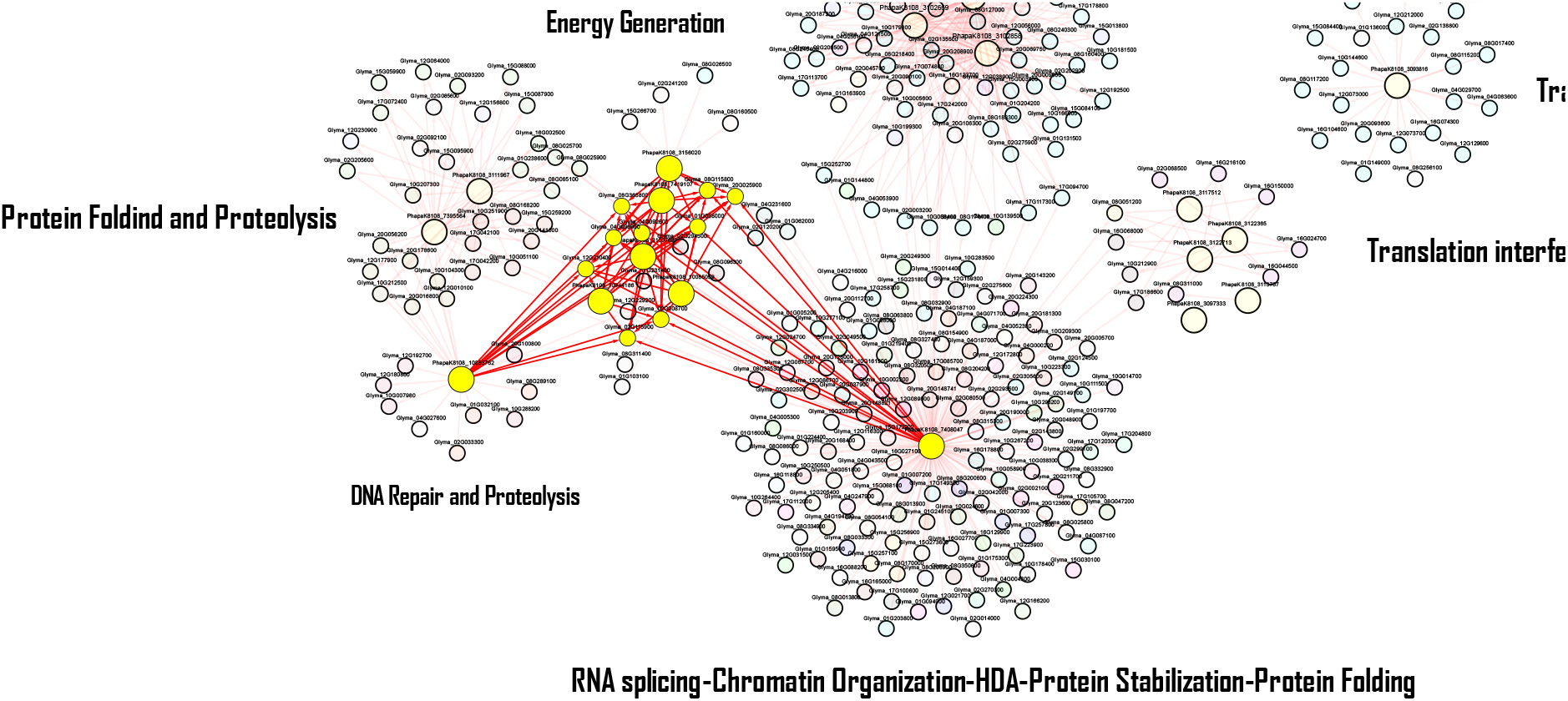
Highlighted, larger nodes represent the effectors; the smaller ones, the SGF14f/14-3-3 target proteins.

Members of this family have been associated with interaction with MYB-type transcription factors in soybean, which in turn modulates nitrogen fixation functions through the CHS8 gene. The study suggests that the interaction between MYB and 14-3-3 controls the intracellular localization of TF (Li;Chen; Dhaubhadel, 2012). Taking into account not only the influence of the balanced nutritional aspect and resistance of plants, since nitrogen fixation is an important process for legumes, MYB is directly involved with plant immunity (Yu et al., 2023). MYB15 in A. thaliana regulates lignin biosynthesis in light of ETI. Lignin is crucial for plant defense, acting as a physical barrier against pathogens. In MYB15 mutant plants, lignin production and disease resistance are compromised by demonstrating that this TF regulates key genes of this biosynthesis (PAL, C4H, 4CL, HCT, C3′H, COMT and CAD), demonstrating the central role of this factor in plant defense (Kim et al., 2020). This suggests that the aforementioned effectors interacting with 14-3-3 proteins, preventing MYB-14-3-3 association, may indirectly influence components of the soybean defense system.

Within the module, another effector was assigned to function as a serine proteinase (Hengst et al., 2001). PEBP may be linked to the pathogen’s adherence to the host cell, which therefore contributes to infection. It has already been reported that these proteins help regulate proteostasis, lipid/phospholipid metabolism, and also inhibit the kinase cascade (Känel et al., 2022; Bernier et al., 1986;Pyo et al., 2018). This last aspect, although little studied, would be the action of this effector as a virulence component. In Drosophila PPIs interactions included Chaperones, ubikinases, phosphorylation and stress responses (Känel et al., 2022). Showing that her immune system is mediated by these proteins and provides protection during infections, since the aforementioned gene was upregulated (Vierstraete et al., 2004; Levy et al., 2004; Reumer et al., 2009). Taking these aspects into account, and assuming that this is an effector secreted by the pathogen, a possible explanation for its form of action would be to mimic the host’s PEBPs with the aim of manipulating, for example, the folding of proteins via Chaperones or interfering with the kinase cascade.

Another important target of the effectors was Pyruvate dehydrogenase E1, which is part of the enzymatic complex ultimately responsible for carbon entry into the Krebs cycle and conversion of pyruvate into acetyl-CoA (Sgrignani et al., 2018), in addition to NADH used in the biosynthesis of fatty acids (FA) (Tovar-Méndez; Miernyk; Randall, 2003). As we know FA are lipid structures that constitute the cell wall and participate in the basal defense of plants, more than that, some FAs containing 16 and 18 carbons modulate the response against pathogens, as well as PTI and signaling at the cuticle level (Kachroo; Kachroo, 2009). Therefore, given these important points of action, this molecule becomes an important phytopathogenic target.

Both the module assigned to“Protein folding and Proteolysis” and “DNA Repair and Proteolysis”, despite being involved in similar processes, have shown that different groups of proteins are targets of different effectors. In the first module, two carboxylases represented respectively by PhapaK8108_3111967 (carboxypeptidase S1) and PhapaK8108_7395564 (carboxypeptidase Y) made contact mainly with target proteins of the HSP70, polyubiquitin and Carboxypeptidase types. While the DNA repair module, the only candidate effector PhapaK8108_10863762 (Acid protease) presented PPIs for Helicase-like transcription factor (CHR28), AP-1 complex subunit gamma and Cysteine proteinase. Figure 8 highlights these modules. CHR28 is a transcription factor linked to DNA resistance to stress-induced damage in the replication process. It gives stability to the replication fork (Seelinger; Otterlei, 2020). S1-type carboxypeptidase effectors catalyze the hydrolysis of both peptide bonds in proteins and peptides (Deiteren et al., 2007). These enzymes are part of the secretome of many pathogens such as fungi and bacteria and have functions mainly for the survival of the microbe in the host environment and virulence and in essence they are protein inhibitors (Yang et al., 2015; Gomes; Espósito; Baracat-Pereira, 2023). Fungi, for example, secrete peptidases to facilitate invasion (Alby; Schaefer; Bennett, 2009).

**Figure 8.**
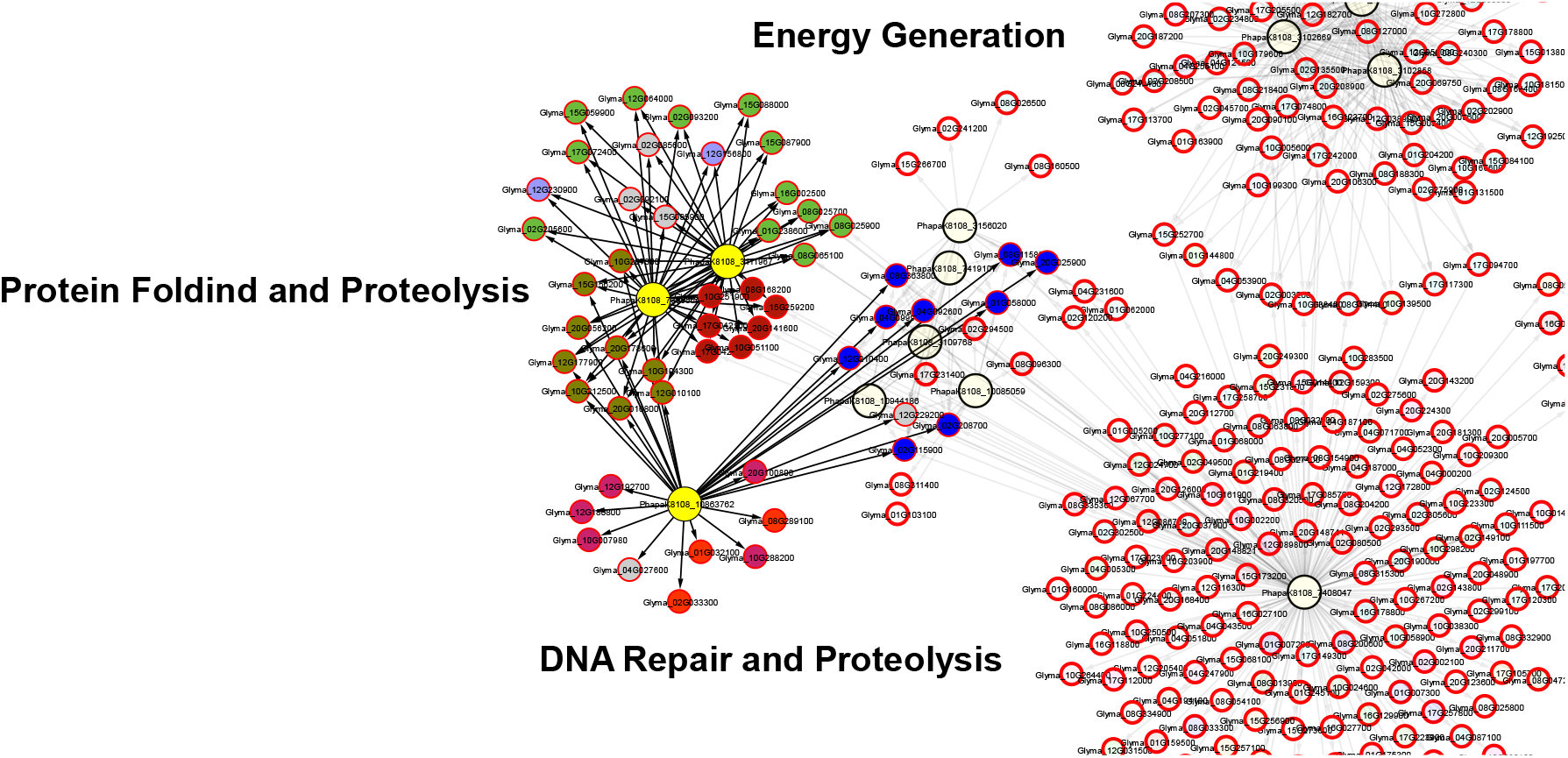
Highlight of the Protein folding/proteolysis and DNA repair/Proteolysis modules. In yellow the effectors; in color, the different classes of proteins.

In general, the repertoire is vast and the types of enzymes secreted target specific substrates (Sriranganadane et al., 2011). In this module, two were identified and 11 were spread across the various modules, with those discussed here interacting with the largest number of targets (33 each). It is interesting to note that within the “Protein folding” module, both carboxypeptidases interact with target proteins from the same group, that is, nine host carboxypeptidases (fig. 7, darker shade of green). This suggests a potential “mimicry” strategy on the part of the pathogen with the aim of evading the surveillance system.

Returning the focus to the DNA repair module, two classes of proteins attract attention. It is DNA repair protein RAD16 and ATP binding hydrolase that interact with the Acid protease effector (PhapaK8108_10863762). Secreted acidic proteases are generally virulence factors in the initial stages of infection, degrading or inhibiting proteins such as polygalactoronase inhibitors (PGIP) (Poussereau et al., 2001; Urbanek; Yirdaw, 1994) and also by loosening the cells that make up the cell wall. (Duby; Boutry, 2009). Secreted proteases are used both to modulate the expression of some genes for successful invasion, as well as a hiding mechanism from the host’s surveillance system. Another aspect that can be attributed is that they also help to deactivate this system (Figaj et al., 2019). A well-studied case of this action is in *P. syringae* pv. tomato in which the bacteria evade the detection system, that is, the specific recognition by the host PRRs of the effectors (flagellin). The mechanism to circumvent the system is as follows, the bacteria secrete a type of protease in the apoplast which in turn degrades the bacterial flagellin itself, which is a powerful inducer of the host innate system, as a result, the plant does not recognize the structure that is played. in half as a bait (Bardoel et al., 2011; Komoriya et al., 1999). Thus, as can be seen, proteases are extremely important in the infection process and become indispensable to the arsenal of *P. pachyrhizi*.

Thinking about the likely function of the Acid protease identified here interacting with the mentioned targets, it could be argued that the inhibition of the host protein (Glyma_10G007980) with conserved helicase domains, as well as RAD16, Cysteine proteinase and AP-1 gamma complex subunit would be its main goal. RAD16, which is part of the repair system, within the context of nucleotide excision, is essential to maintain integrity. It plays important roles in the repair of genes that are being encoded or takes part in the repair mainly of repressed genes (Li et al., 2008), this process is dependent on ATP (Guzder et al., 1998) and may explain why effector of this module (DNA Repair, fig. 3,7) possibly act in some way by inhibiting this chain. The HPI network identified the target protein ATP binding-hydrolase interacting with the effector (PhapaK8108_10863762), and as it is an acidic protease with inhibitory action, it can act in this direction. This would indicate that not only does it target components of the repair machinery, but also subcomponents such as enzymes that transfer ATP to the system to activate it. ATP hydrolase-type molecules, as is the case, normally trigger, through ATP hydrolysis, the translocation of RAD16 along the strand in search of damage or lesions of genes that will be transcribed or are in a repressed state (Guzder et al., 1998). Indicating a cooperative and complex action of phytopathogenic effectors to influence host destabilization, the underlying effects of this on the negative regulation of the immune system are still unclear. And this inhibition of ATPase by pathogens has already been confirmed (Sondergaard et al., 2004). Although the mechanical form of action of some protease-type effectors is unclear, they act by destabilizing the function of ATPases, possible pathways that may lie in analogous functions of toxins used to suppress this system. Reports point to an indirect action on membrane fluidity that allows the activity of proton pumps, while other studies suggest that these toxins can disrupt the function of the host membrane by forming pores that dissipate the electrochemical gradient of protons across the membrane (Simon-Plas et al., 1996; Goudet et al., 2000).

MAPK is one of the main downstream mechanisms in response to pathogenic attack, since the activation of the cascade and consequently of the responsive genes, mediated by plant hormones, initially occurs through this pathway (Sun; Zhang, 2022). Proteins with kinase domains were the preferred targets for effectors in our study. Looking at Figure 3, we identified two protein-rich modules from the kinase superfamilies (nodes in light blue). Despite this richness mentioned, the effectors participate in different biological processes (“Endocytosis and blue light signaling pathway” and “Signal Transduction”). There was a separation, for example, with the effector with an uncharacterized hypothetical protein annotation for the first module and a glucanase for the second. This demonstrates and reflects the complexity and diversity of functions of this super family (Boutrot; Zipfel, 2017; Klymiuk et al., 2021). The family with kinase domains, despite their structural differences and spatially distinct cellular locations, their activities diverge greatly. They are linked to signaling, Ca2+ influx, activation of mitogen-activated protein kinase (MAPK), extracellular production of reactive oxygen species, transcriptional reprogramming and formation of physical barriers for cell fortification (Zhang et al., 2020). Considering this aspect, the action of both the effector PhapaK8108_11631953 (hypothetical protein) and PhapaK8108_3093816 (Glucanase) is at the ETI level and, therefore, already within the scope of gene-by-gene and species-specific theory. Glucanases are enzymes used mainly to degrade cell walls and act in programmed cell death (Ökmen, Bachmann; de Wit, 2019), therefore at the level of PTI or basal defense (Zhu et al., 2017). Within the context of the action of glucanases on kinases, since we identified 19 host proteins containing this domain in this module, it is still not very clear. However, there are reports of effectors of this type in *Botrytis cinerea* that are located in extracellular spaces and can act by interacting with membrane proteins with kinase domains and then activating the signaling system via BAK1 (Zhu et al., 2017). This suggests that one of the possible actions of the pathogenic effector is to manipulate this system in order to reduce or completely mask its presence in the face of the host’s sophisticated system.

And last but not least, the module attributed to action on carbohydrate metabolism that presented two potential effectors with two target host proteins. Two Peptidoglycan deacetylase effectors (PhapaK8108_9190215 and PhapaK8108_3166594) were identified and both are from the Carbohydrate esterases family, the latter being corroborated by the analysis of Carbo-active effectors (fig. 1). The two interacted with the host proteins Membrane-bound O-acyltransferase (MBOAT), which is part of an important family of proteins dedicated to the catalysis of fatty acyl groups in lipids and proteins. Several types of secreted substances have already been reported to be substrates for MBOAT action (Buglino; Resh, 2012). Membranes are not static, they are dynamic interfaces used for signal transmission, traffic and interactions. Modifications, such as the addition or removal of lipids, not only allow the binding of proteins to the membrane, but also modify their intrinsic function (Hemsley, 2014). Thus, it is reasonable to think that such modifications promoted by certain effectors can destabilize them to avoid triggering the signaling system (Zhang et al., 2022). In addition to this, it is known that if membranes are embedded with receptor proteins (PRRs) that recognize the molecular patterns of microbes (PAMPs), changes or imbalances in their components can influence the speed or even specific protein-protein recognition (Li; Wu, 2021; Monaghan; Zipfel, 2012). In fact, taking into account the aspects of MBOAT having secreted proteins as substrates, one could think that the effectors of *P. pachyrhizi* could go unnoticed, if they are similar to those of the host. The pathogen would usurp the host’s own machinery to add modifiers that would help cross the membrane. However, the two effectors, as indicated by their carbohydrate activity, would be modifying agents of the host MBOATs or even modifications of the cell wall itself (Moynihan; Sychantha; Clarke, 2014) so as not to be recognized by the immune system and subsequent passage through the membrane with the help of the host MBOAT. It should be noted that this is merely intuitive considering the aspects discussed here, since no direct relationship was found between MBOAT and carbo-active proteins of the esterase type.

In general, it is noted that the HPI network presented in this article showed that there is a great complexity between the processes, as well as the proteins that are targets of *P. pachyrhizi*. Despite major advances in bioinformatics, such as the application of deep learning to improve predictions, many gaps and inconsistencies are still observed. However, it does not invalidate the general scope of the knowledge developed. Future studies that aim to characterize, through silencing, the potential effectors found here will be able to prove their activities within the infection process.

## 4.0 CONCLUSIONS

Thorough analysis of the interactions between fungal effectors and the soybean proteome revealed crucial insights into the molecular mechanisms of plant-pathogen interactions. Using the PredHPI tool, 922 unique protein-protein interactions were predicted between the 185 effectors of the fungus and the soybean proteome, demonstrating the effectiveness of this computational approach in generating intraspecific protein-protein interaction networks and which can be followed for other pathosystems.

Pathogen effectors, especially Lysophospholipases (LysoPLs), Subtilisin Proteases and Glycoside Hydrolases, have been identified as potential effectors within the arsenal used by *P. pachyrhizi*, with multiple effectors in each class. The effector PhapaK8108_7408047 (Cyclophilin-like) stood out for having the largest number of target proteins, suggesting its importance in the virulence of the pathogen and its interactions with various cellular processes, such as RNA splicing, chromatin organization and protein stabilization.

Other effectors showed significant interactions in modules related to vesicle trafficking, protein translation and energy metabolism, indicating pathogen strategies to manipulate essential cellular processes and subvert the host immune response. Furthermore, effectors also interacted with proteins in important signaling pathways, such as the MAPK pathway, highlighting the complexity of plant-pathogen interactions and the pathogen’s ability to modulate host cellular responses.

These findings provide a solid basis for developing plant disease control strategies and for a deeper understanding of host-pathogen interactions, potentially paving the way for more effective interventions in the future. More targeted approaches in the future, such as silencing these key effectors, could provide solid insights into effector biology.

## Supporting information

supplemental Blast2Go_results

## Notes

### Competing Interest Statement

The authors have declared no competing interest.

